# Intrinsic K-Ras dynamics: A novel molecular dynamics data analysis method shows causality between residue pairs

**DOI:** 10.1101/067496

**Authors:** Sezen Vatansever, Zeynep H. Gümüş, Burak Erman

## Abstract

While mutant K-Ras is an important therapeutic target for human cancers, there are still no drugs that directly target it. Recent promising studies emphasize the significance of dynamics data to selectively target its active/inactive states. However, despite tremendous information on K-Ras, the direction of information flow in the allosteric regulation of its dynamics has not yet been elucidated. Here, we present a novel approach that identifies causality in correlated motions of proteins and apply it to K-Ras dynamics. Specifically, we analyze molecular dynamics simulations data and comprehensively investigate nucleotide-dependent intrinsic K-Ras activity. We show that GTP binding leads to characteristic residue correlations with relatively long decay times by stabilizing K-Ras motions. Furthermore, we identify for the first time driver-follower relationships of correlated motions in the regulation of K-Ras activity. Our results can be utilized for directly targeting mutant K-Ras in future studies.

## Introduction

K-Ras is a small GTP-binding protein pivotal in cellular signaling. Somatic K-Ras mutations are among the most common activating cancer lesions, driving 71% of pancreas, 35% of colon and 17% of lung cancers (Slebos et al., 1990, Stephen et al., 2014, Forbes et al., 2015). Signaling through K-Ras is dependent on the bound nucleotide, where the GTP-bound state is active (on) while the GDP-bound state is inactive (off). In GTP-bound K-Ras, P-loop (residues 10-17), switch I (SI, residues 25-40) and switch II (SII, residues 60-74) regions make up the active site whose well-ordered conformations allow effector protein binding for protein signaling (Figure 1). However, oncogenic gain-of-function mutations impair GTP hydrolysis and freeze K-Ras in its active state (Vetter and Wittinghofer, 2001), causing uncontrollable cellular growth and evasion of apoptotic signals (Downward, 2003, Chen et al., 2013). Tumors driven by oncogenic K-Ras are often resistant to standard therapies and result in poor outcomes; they are also excluded from treatment with other targeted therapies, making mutant K-Ras an extremely high priority target in cancer treatment (Pao et al., 2005, Lievre et al., 2006). However, there are still no drugs in the clinic today that directly target mutant K-Ras.

**Figure 1.**
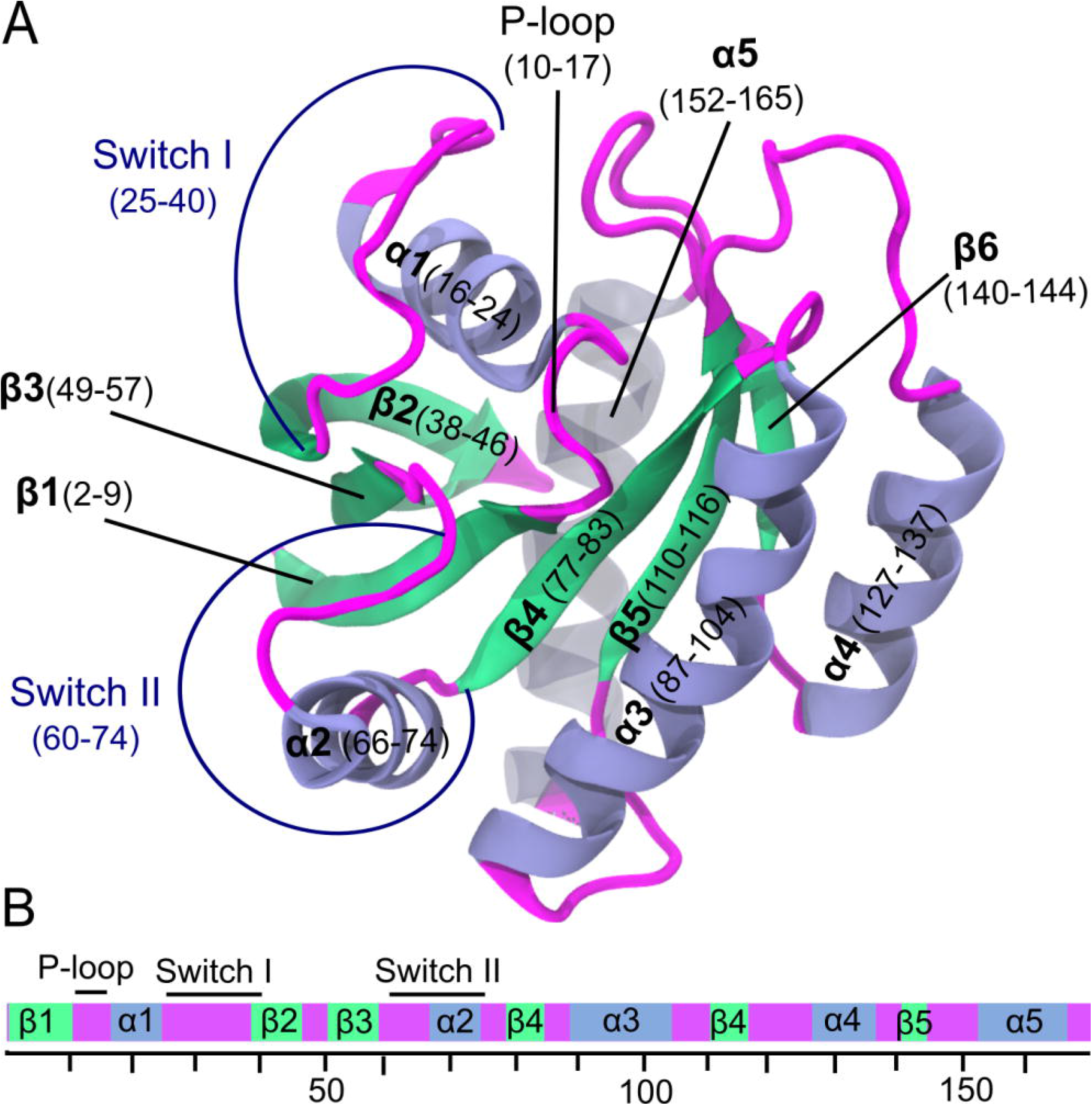
Crystal structure of wild-type K-Ras protein in GTP-bound state (PDB: 4OBE). A) K-Ras structure ribbon representation with secondary structures in blue for α-helices and green for β-sheets. B) Schematic of K-Ras sequences (residues 1-169). Functional regions are in same color used in K-Ras structure in A.

Part of the challenge in oncogenic K-Ras inhibitor design has been due to structure analyses that suggest a lack of well-defined druggable sites on its surface (Zhang and Cheong, 2016). However, studies that have utilized protein dynamics data such as NMR and mass spectrometry have identified binding pockets on specific K-Ras oncogenic mutants and have attempted to stabilize their on or off conformational states (Taveras et al., 1997, Ostrem et al., 2013, Lim et al., 2014, Lito et al., 2016). Accumulating studies suggest that K-Ras proteins are in dynamic and flexible states and their distinct characteristics cannot be identified by structural studies alone (Patricelli et al., 2016, Ostrem et al., 2013, Lim et al., 2014, Singh et al., 2015, Grant et al., 2011b, Leshchiner et al., 2015, Lito et al., 2016). K-Ras dynamic activities in different conformational states, which can also change due to allosteric interactions between protein residues, also need to be quantified (Motlagh et al., 2014).

However, we still do not clearly understand the intra-molecular allosteric networks between distant sites on K-Ras (Marcus and Mattos, 2015). While such allosteric interaction sites have recently been discovered in its catalytic domain (Buhrman et al., 2010, Abankwa et al., 2010, Kearney et al., 2014), they remain largely understudied (Marcus and Mattos, 2015). Understanding allosteric interactions can present novel opportunities for small molecule targeting of mutant K-Ras conformations while sparing those of wild-type K-Ras, which first requires a deeper understanding of the intrinsic K-Ras dynamics.

In allosteric regulation of protein dynamics, correlated motions between protein residues are essential (Goodey and Benkovic, 2008, Kern and Zuiderweg, 2003, Wand, 2001). These motions enable the transfer of fluctuation information through the allosteric network (Kamberaj and van der Vaart, 2009), which inherently involves directionality, or “causality” of events (Guarnera and Berezovsky, 2016). If the motions of two residues are correlated, it would be extremely valuable to identify whether the motions of one residue drive the motions of the other. However, while correlation calculations indicate interaction (which is necessary for allosteric transitions), they are symmetric and do not reveal the direction of information flow. To understand the directionality (or causality) between the motions of two residues, we need to incorporate time-delayed correlation estimations in our analyses (Schreiber, 2000, Guarnera and Berezovsky, 2016).

Here, we introduce a novel method that enables identification of causality between all residue pairs of a protein using time-delayed correlation information. For this purpose, we first record residue fluctuations calculated at every time step of a molecular dynamics (MD) simulation as a time series, and then calculate the time-delayed correlation of a residue pair as the correlation between two time series where one is shifted in time relative to the other. We conclude that the fluctuations of a residue control and modify the fluctuations of the delayed one if we observe a significant correlation between the fluctuations of two residues with a positive time lag. This is a completely novel approach to the analysis of structure-function relations, as well as to understanding K-Ras dynamics. We demonstrate the simplicity of computing time-correlation functions in studying protein dynamics by applying the method to study K-Ras. We specifically focus on K-Ras because of its clinical importance and the availability of a large body of data which enable the validation of our theoretical predictions. Note that while correlations between the fluctuations of residue pairs have already been shown in several Ras protein studies (Grant et al., 2009b, Lukman et al., 2010, Kapoor and Travesset, 2015), despite the significantly large body of literature on K-Ras, there has been surprisingly little attention on the role of causality (or directionality) in correlation dynamics of K-Ras conformations.

In summary, we present a comprehensive study of intrinsic K-Ras dynamics, including detailed analyses of causality between the motions of its residues. We first provide detailed, quantitative descriptions of both active and inactive K-Ras from extensive MD simulations. We use a statistical thermodynamics interpretation of fluctuation correlations to quantify K-Ras ‘stiffening’ upon activation. Using stiffness calculations jointly with measurements of reduced relative fluctuations, we define protein stability and show that K-Ras is more stable in active conformation. To investigate time-dependent characteristics of correlated motions, we map and compare correlated motion patterns of active and inactive K-Ras, then discuss the differences of decay times of correlations between the two K-Ras forms in detail. Our results show that inactive K-Ras is marked by a pronounced decrease in correlated motions of residues for shorter periods, while active K-Ras correlations have longer decay times. We analyze the ensuing events at the atomic scale. Finally, to enable a deeper understanding of K-Ras dynamics, we introduce the first causality calculations for K-Ras and identify specific driver and follower residues during protein simulations.

## Results and Discussion

### Comparison of stiffness changes in active and inactive K-Ras

#### GTP binding increases K-Ras stiffness

To understand how nucleotide binding effects K-Ras dynamics, we quantified changes in its ‘stiffness’ – a metric that inversely correlates with residue pair fluctuations - upon GTP vs. GDP binding. For this purpose, we represented the interaction between two fluctuating residue pairs (*i* and *j*) as a spring with a constant *k_ij_*, based on a statistical thermodynamics interpretation we previously developed (Erman, 2015). Plotting this spring constant for every residue pair in both GTP-(Figure 2A) and GDP-bound (Figure 2B) K-Ras, we observe strong coordination in the fluctuations of GTP phosphate groups with those of K-Ras (Figure 2A).

**Figure 2.**
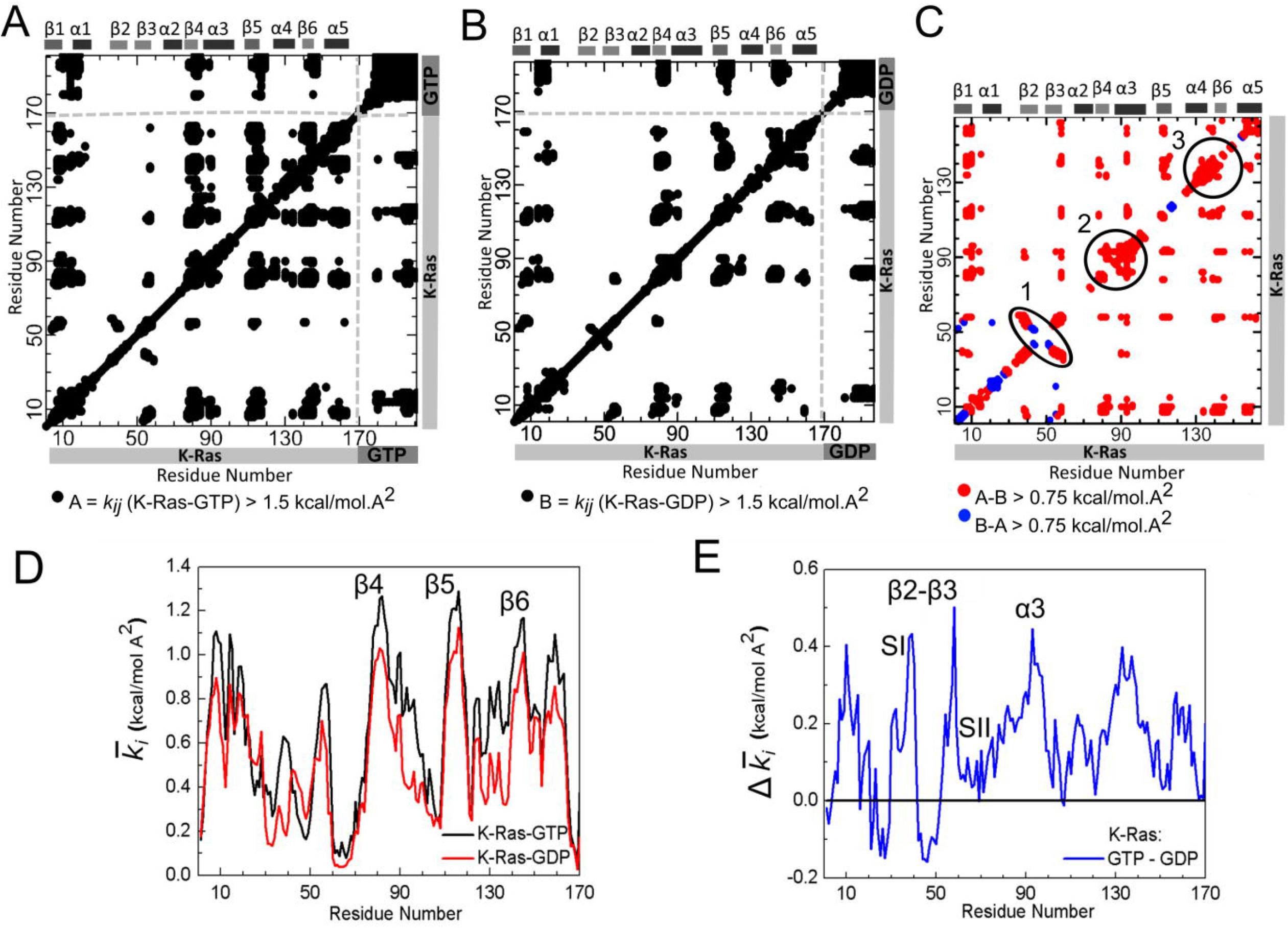
Stiffness results for GTP- and GDP-bound K-Ras and their difference. In panels A and B, both axes marks 1-169 represent the residue Cα atoms of K-Ras and marks 170-on represent GDP and GTP nucleotide heavy atoms, respectively with *k_ij_* >1.5 kcal/mol A^2^. (A) *k_ij_* for K-Ras-GTP. Atoms 170-181 are the γ, β, α-phosphate groups and 182-201 are the guanine atoms of GTP. (B) *k_ij_* for K-Ras-GDP. Atoms 170-178 are the (β and α phosphate groups and 178-197 are the guanine atoms of GDP. (C) Difference between active and inactive K-Ras *k_ij_* values. Red regions are stiffer in K-Ras-GTP (*k_ij_* values of K-Ras-GTP are greater than K-Ras-GDP at least 0.75 kcal/mol·A^2^) and blue regions are stiffer in K-Ras-GDP (*k_ij_* values of K-Ras-GDP are greater than K-Ras-GTP at least 0.75 kcal/mol·A^2^). (D) Mean spring constants 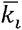 for the GTP and GDP bound states. (E) Mean spring constant differences 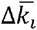 for the GTP and GDP bound states. Positive values correspond to larger mean stiffness in K-Ras-GTP.

To zoom in on and directly compare the effects of nucleotide binding on K-Ras stiffness, we simply calculated the differences in spring constant values between GTP- and GDP- bound K-Ras. In Figure 2C, red dots indicate that the differences are largely due to the stiffening effects of GTP-binding on residue pair fluctuations. Notice that Regions 1-3 in Figure 2C that correspond to secondary structures show significant increase in *k_ij_* when GTP-bound. Furthermore, Region 1 corresponds to strong coordination of β2 and β3 motions, while Regions 2 and 3 correspond to increased stiffness of β4-α3 and α4.

#### Nucleotide binding affects spring constant of α2 (SII)

We next investigated the effects of nucleotide binding on the spring constant of α2 (SII), because previous studies have shown that stiffness increases when SII refolds into an α-helical conformation through GTP binding (Noe et al., 2005). We calculated the spring constants of the two terminal residues of α2 (A66 and T74), which were 0.10 kcal/mol·A^2^ (69.91 pN/nm) for active and 0.04 kcal/mol·A^2^ (27.78 pN/nm) for inactive K-Ras. Previous studies have utilized various experimental methods that have all led to spring constants within ~0.09-1.15 kcal/mol·A^2^ (60–80 pN/nm) for helices (Adamovic et al., 2008, Howard, 2001). Our results for both forms of K-Ras are in the same order of magnitude. Note that for active K-Ras the α2 spring constant is equalent to the characteristic spring constant of α-helices, while it is lower in inactive form. Hence, our results validate and quantify earlier, qualitative observations of Noe *et. Al*. (Noe et al., 2005) that the α2 spring constant reaches to the level of an α-helix spring constant during GTP binding.

#### Overall spring constant is higher in active complex

To estimate global changes in stiffness in response to nucleotide binding, we calculated the overall spring constants *k_overall_ (details in Methods)* of nucleotide-K-Ras complexes, which were 0.70 kcal/mol·A^2^ (481.75 pN/nm) for GTP-bound, and 0.55 kcal/mol·A^2^ (385.12 pN/nm) for GDP-bound K-Ras. Both are of the same order of magnitude with an experimental study for another protein, myoglobin, which has an overall spring constant of ~300 pN/m (Rico et al., 2013, Zaccai, 2000) pointing to an order of magnitude agreement of overall stiffnesses of proteins in general. In conclusion, GTP-binding makes K-Ras more rigid.

#### Secondary structure motions show the strongest coordination with the rest of the protein

Quantifying the spring constant based on fluctuations allows for analyzing how, analogous to a virtual spring, the fluctuations of a specific residue are coupled with fluctuations of rest of the protein. To discover residues whose fluctuations are in strong coordination with K-Ras fluctuations and how they change between the two states, we compared the mean spring constant 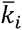 of each residue *i*, for both active and inactive K-Ras (Figure 2D) as described in *Methods*. A large 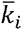 value indicates that the motions of residue *i* are stiffly coupled with the motions of the protein; while a small 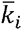 value indicates that the motions of the *i*th residue and the protein are flexibly coupled. For simplicity, we categorized the significant mean spring constant 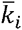 values as highest, high and smallest (For details please see Table S1). In both states, the highest 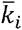 values are of β-strand residues β4, β5 and β6, showing the strongest coordination of their motions with K-Ras motions. Next, high 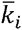 values of β1, P-loop and α5 residues indicate that their fluctuations are also strongly coupled with those of the protein. On the other hand, the smallest 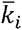 values belong to SII region in active and SI and SII regions in inactive K-Ras which show that their residue fluctuations are not correlated with the rest of the protein (Table S2). Since we have defined the stiffness metric as a signifier of a decrease in residue fluctuations, we provide a second line of proof that increased stiffness stabilizes dynamic fluctuations in both forms of K-Ras by using Root Mean Square Fluctuation (RMSF) graph (Figure S1). Clearly, the residues with the smallest mean spring constant 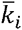 values from Figure 2D have the highest RMSF values in Figure S1 and vice versa.

As indicated in previous studies where NMR and Atomic Force Microscopy were used, protein stiffness depends on secondary structure (Zaccai, 2000, Rico et al., 2013), where loops contribute to structural flexibility and show large fluctuations, while β-strands and α-helices provide mechanical stability and show small fluctuations (Rico et al., 2013). Our K-Ras results are consistent with these general observations. In addition, we observe stiff coupling of the fluctuations of the P-loop and the protein. This observation is important since P-loop is the phosphate binding site of K-Ras and connects β1 and α1 (Figure 1). Although loops are often flexible regions of proteins and show higher fluctuations, in K-Ras, motions of P-loop residues are stiffly coupled to those of the protein, especially in active state (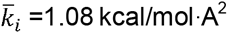 for K-Ras-GTP, 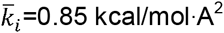 for K-Ras-GDP).

#### The mean spring constant values of residues in β2, β3, α3 and switch regions – especially SI- are higher in active K-Ras than in inactive K-Ras

Finally, we calculated mean spring constant differences between active and inactive K-Ras, 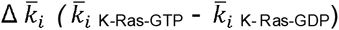. Figure 2E shows that the fluctuations of β2 and β3 terminal (D38 and D57) and α3 center (D92-I93) residues are in stronger coordination with those of active K-Ras (vs. inactive K-Ras) 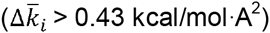. Our results also indicate that although residues of switch regions have the smallest 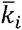 values in both forms, some of their 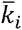 values increase significantly in active form. In Figure 2E, 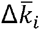 ranges between 0.20–0.36 kcal/mol·A^2^ for residues in SI (D30-R41) and 0.02-0.19 kcal/mol·A^2^ for residues in SII (G60-T74). These 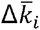 values show stiffer coupling of the motions of GTP-K-Ras with the motions of switch residues, especially SI (vs GDP-K-Ras). This result is important as SI includes the binding site to effector proteins which only bind to GTP-bound K-Ras when SI flexibilty is reduced (Spoerner et al., 2010). Earlier studies that used NMR spectra and RMSF calculation also support our results that GTP binding reduces the flexibility of both SI and SII, especially SI (Shima et al., 2010, Kapoor and Travesset, 2015). Our results improve on this information by showing that fluctuations of switch regions–notably SI- are more stiffly coupled with K-Ras-GTP fluctuations (Figure 2E).

### Comparison of residue pair correlations for active and inactive K-Ras

To identify if the fluctuations of one residue are related to fluctuations of another residue, we calculated the correlations of all residue-residue pairs in both GTP- vs GDP bound K-Ras complexes. As expected, cross-correlation coefficient maps of K-Ras-GTP (Figure 3A) and K-Ras GDP (Figure 3B) exhibit different correlation characteristics. The most remarkable differences between Figure 3A and 3B belong to two parts: (i) the correlation of α1-S1 with L10-α5 and (ii) the correlations between β2 and β3. Positive correlation patterns within these two parts are evident in K-Ras-GTP simulations, but absent in K-Ras GDP simulations. To provide comprehensive information on nucleotide-dependent K-Ras dynamics, we present these two remarkable results from correlation analyses (Figure 3) as well as sources of correlated motions (i.e. H-bonds) together in the following sections.

**Figure 3.**
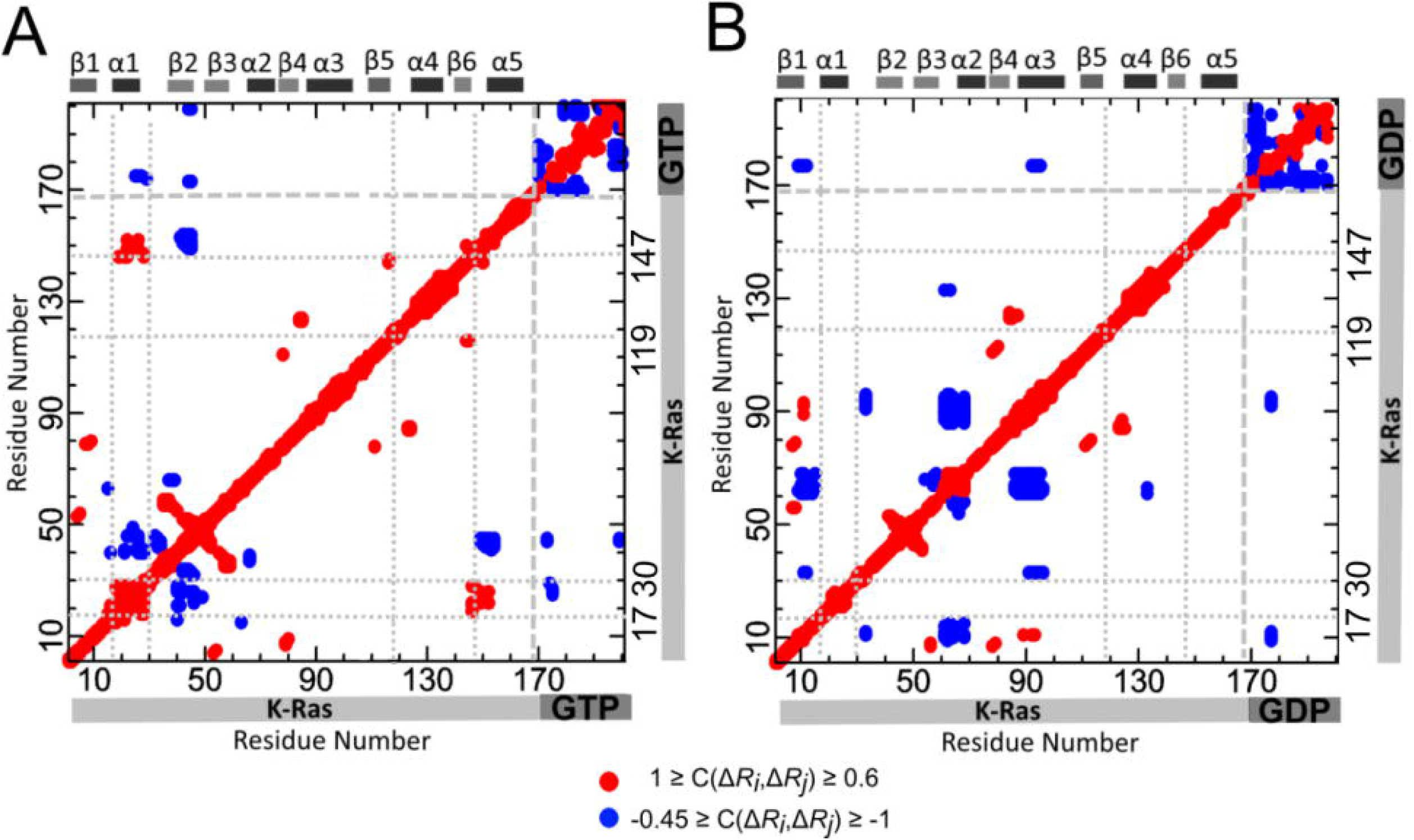
Cross-correlation coefficient maps for GTP and GDP bound states. Red dots show positive correlations (1 ≥ C(Δ*R_i_*, Δ*R_j_*) ≥ 0.6) and blue dots show negative correlations (-0.45 ≥ C(Δ*R_i_*, Δ*R_j_*) ≥ -1). Residues indexed 1-169 belong to K-Ras. (A) Correlated fluctuations of K-Ras-GTP. Indices between 170-201 refer to GTP heavy atoms (182-201 are guanine atoms). (B) Correlated fluctuations of K-Ras-GDP. Indices between 170-197 refer to GDP heavy atoms (178-197 are guanine atoms).

#### The correlation of α1-S1 with L10-α5 in active K-Ras motions is due to three specific H-bonds

MD simulations show that the correlation between α1-S1and L10-α5 in the active form results from GTP binding to active site residues, which also form specific H bonds with other K-Ras residues and water. Based on the average number of H-bonds each residue forms throughout the simulation, we estimated that the nucleotides remain bound to active site residues S17, D30, D119 and K147, and that GTP-binding (vs. GDP) is more stable for S17 and D30 (Table S3). Furthermore, correlated motions of α1-S1 and L10-α5 in GTP-bound K-Ras originate specifically from three H-bonds: (i) A146-Q22, (ii) D30-GTP, (iii) D30-a water molecule. We observed a sustained H-bond between A146-Q22 during active but not in inactive complex simulation. This suggests that A146-Q22 interaction causes a strong relationship between L10α5 (A146-D154) and α1 (L19-I24) in active K-Ras (Figure 3A) with a correlation coefficient of 0.75, and a weak correlation coefficient of 0.28 for inactive K-Ras. At the same time, the active site residue D30 forms an H-bond with the nucleotide in both active and inactive K-Ras, while it also binds to a water molecule only in the active form. However, the H-bond in the active form between D30(O)-GTP(O2A) is more permanent than the H-bond in the inactive form between D30(O)-GDP(O2’).

#### Since H-bond of D30-GTP is permanent, D30-GTP distance is invariant and their fluctuation correlations have longer decay times during K-Ras-GTP simulation

We next combined cross-correlation results with the distance distribution of D30 and nucleotides and quantified the decay times of their correlations during MD simulations. In addition to more permanent binding of D30(O)-GTP(O2A), nucleotide-D30 distance distribution pattern is close to the normal distribution curve with a mean of a smaller value in active K-Ras (Figure 4A), with a correlation coefficient of 0.97. To quantify decay time of this correlation in both complexes, we first defined two “connectivity vectors”, *ΔR_30-GTP_* and *ΔR_30-GDP_*, between D30(O) and nucleotides. As illustrated in Figure 4B, *ΔR_30-GTP_* connects the starting point of fluctuation vector of *ΔR_D30(O)_* to end point of negative *ΔR_GTP(O2A);_ ΔR_30-GDP_* starts from *ΔR_D30(O)_* to negative *ΔR_GDP(O2’)_*. We then calculated time-delayed autocorrelations of each connectivity vector throughout the MD simulations. The autocorrelation plot in Figure 4B summarizes the correlation of connectivity vectors at various time delays, where vector correlation coefficients are plotted with 1 ns delays at a time; slow decay of correlations in active K-Ras is clearly observed. Correlations decay to 1/e in about 3 ns for K-Ras-GDP (red line), vs. to ~10 ns for K-Ras-GTP (black line). One reason for this slow correlation decay is the H-bond, which binds D30 to a water molecule in active K-Ras. The O atom of D30 establishes an H-bond with the nearest water during 28% of the trajectory while it does not make any contact with waters when K-Ras is inactive.

**Figure 4.**
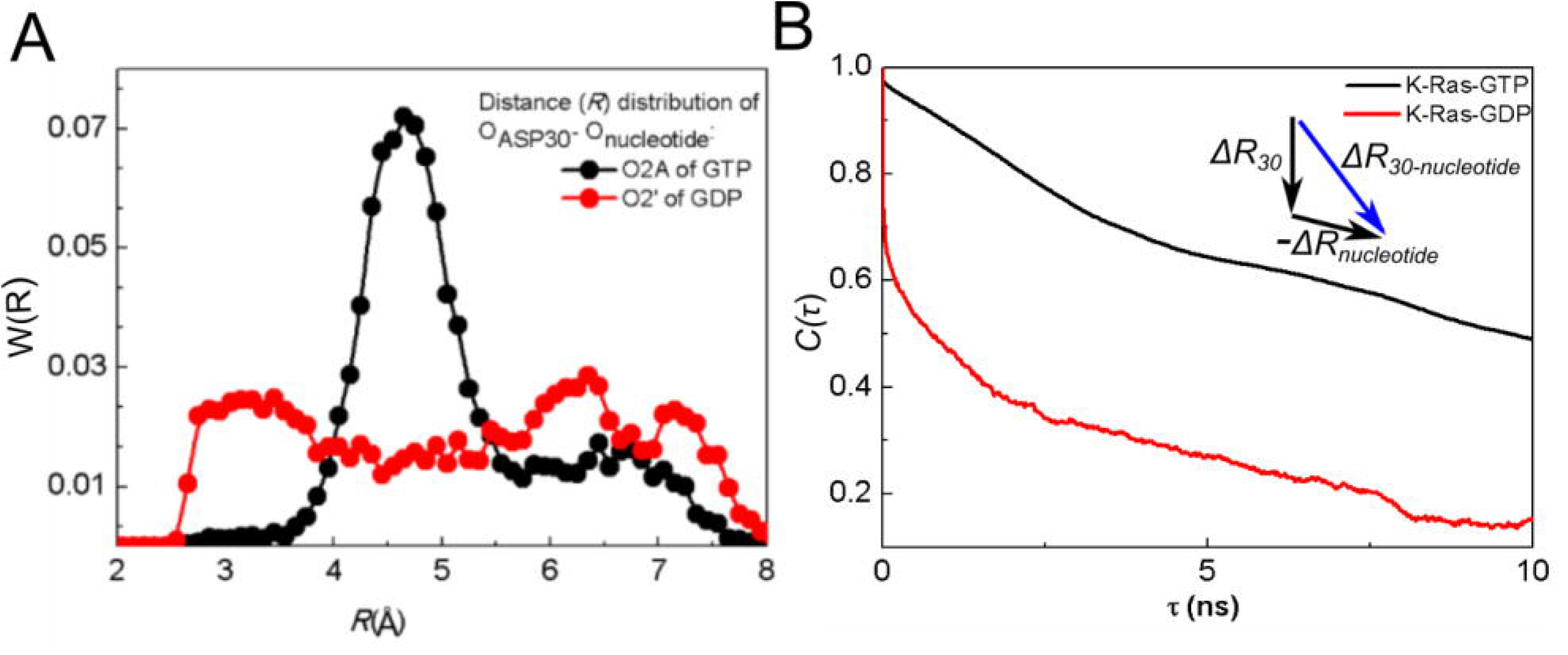
D30-GTP distance is more stable than that of D30-GDP. Fluctuations of D30(O) to GTP(O2A) “connecting vector” are persistently correlated. (A) Distance distribution between D30 and connecting O atoms of GTP (black) and GDP (red) (B)Time delayed autocorrelations for the vector connecting Oxygen atom of D30 to O2A of GTP (black curve) and O2’ of GDP (red curve). X-axis is the time delay (*τ*) and Y axis is the time delayed autocorrelation of the vector for *τ*.

#### A continuous H-bond stabilizes β2-β3 distance and promotes longer decay times for β2-β3 correlations during K-Ras-GTP simulation

β2 and β3 are two parallel β strands located between SI and SII regions (Figure 5A). Due to the presence of a persistent H-bond between R41(β2)-D54(β3) in K-Ras-GTP simulation, the peak value of *R_41-54_* distribution decreases (Figure 5B) and fluctuations of β2 and β3 become correlated (Figure 3A). The time-delayed autocorrelations of the vector *ΔR_38-57_* between their terminal residues D38 and D57 are presented in Figure 5C showing that *ΔR_38-57_* correlation decays much more slowly in active K-Ras.

**Figure 5.**
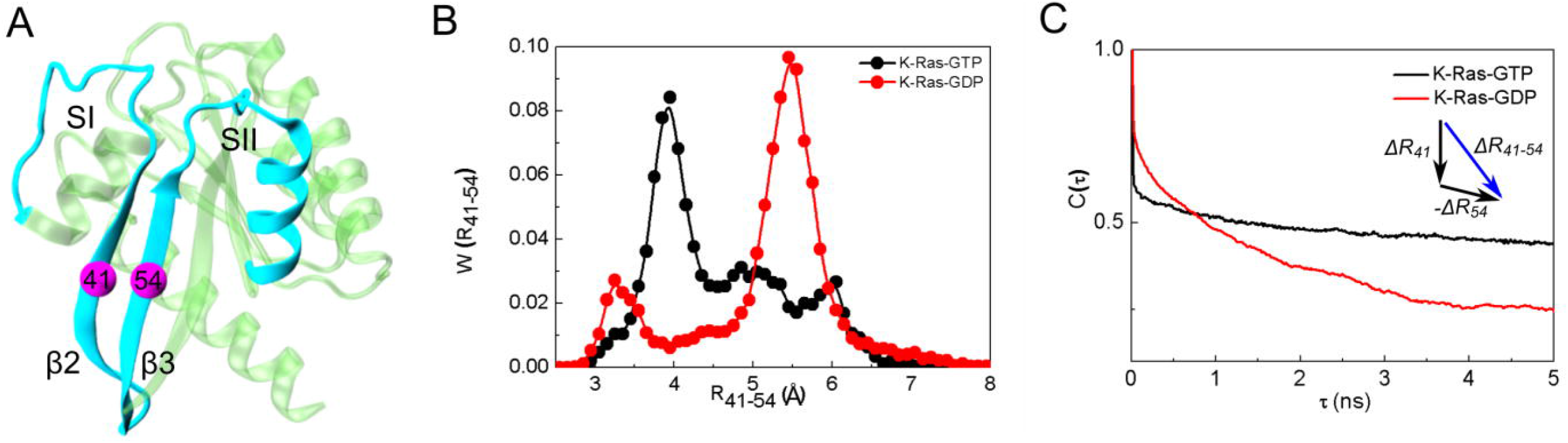
Correlation of β2 and β3 fluctuations is persistent in active K-Ras. (A) Locations of R41 (β2) and D54 (β3) relative to SI & SII. (B) Distance distribution between Cα atoms of R41 and D54 in K-Ras-GTP (black) and K-Ras-GDP (red). Distance values between R41 (β2) and D54 (β3) populate at 3.90 Å during GTP binding whereas these distance values populate at 5.46 Å for GDP-bound K-Ras. (C) Time delayed autocorrelations for the fluctuations of the vector from D38 (Cα) to D57 (Cα).

### Causality of Correlated Motions

Correlated motions of proteins often have a direction or causal relationship (Kamberaj and van der Vaart, 2009). Correlations in the fluctuations of two residues indicate interaction, which is necessary for allosteric transitions. However, this is not sufficient for understanding the dynamic phenomenon completely since these symmetric correlations do not contain information on driver and follower relationships. To deduce causality, time-delayed correlations need to be analyzed. Our trajectory analysis shows that the symmetry assumption on time-delayed correlations does not hold (Callen, 1985) for several residue pairs i.e., time delayed correlations of fluctuations of two atoms are not the same in forward or backwards in time. Our observation is supported by recent work (Kamberaj and van der Vaart, 2009) that identified causality in correlated motions from MD simulations using an information theory measure of transfer entropy. This work, in turn, was built on a study by Schreiber, who introduced the entropy transfer concept for fluctuating environments (Schreiber, 2000). We follow up on these ideas and introduce a new method to dissect time-dependent correlations of all residue pairs of a protein to identify driver and follower residues. For this purpose, we evaluate strong time-delayed (τ=5ns) correlations between residue pairs (defined as < -0.3 or >0.6). The strongest causal relations are as follows (Figure 6):

**Figure 6.**
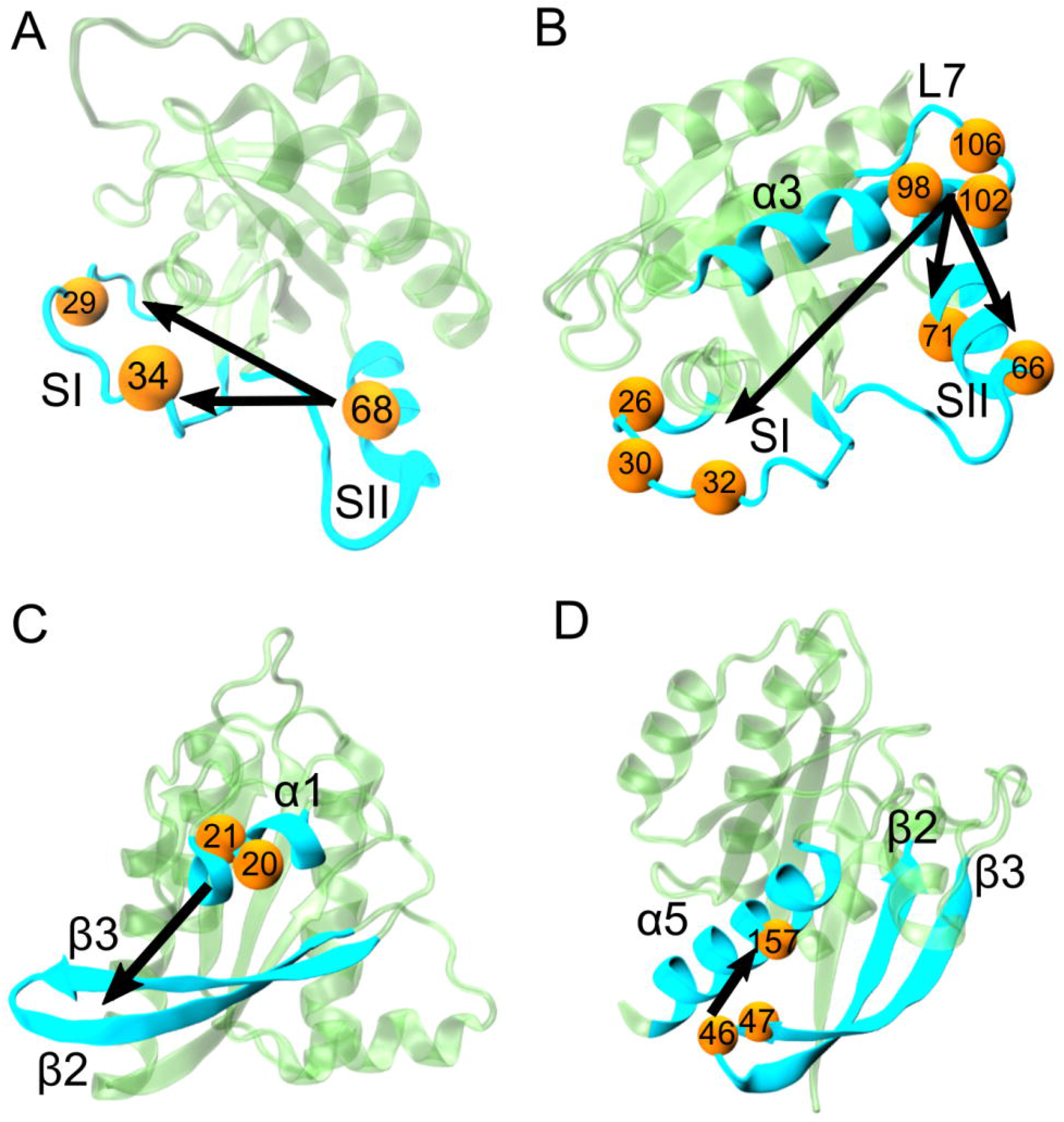
Causality relations in active K-Ras motions. Directionality in causal relationships is illustrated with arrows. Arrows start from driver residues and end at follower residues. Both residue types are represented with yellow spheres and marked with their residue numbers. The secondary structures they belong to are in turquoise. A) R68 (SII) drives V29 and P34(SI). B) E98 and R102 (α3) drive A66 (α2; SII). S106 (L7) drives Y71 (α2; SII). R102 (α3) drives N26 and Y32(SI). S106 (L7) drives D30. C) ILE21-GLN22 (α1) drives β2-β3. D) I46 and D47 (**β2-β3**) drive Y157 (α5).

#### SII motions drive SI in active K-Ras

SI-SII relationship is better understood by examining residues that drive their motions throughout the trajectory. Our causality calculations show that SI is driven by SII (Figure 6A and 7). We present time-delayed correlation plots of R68(SII) with V29(SI) (Figure 7A) and with P34(SI) (Figure 7B) for active K-Ras. Red curve shows that the fluctuations of R68 at time *t* affect the fluctuations of V29 at time *t*+*τ*.

**Figure 7.**
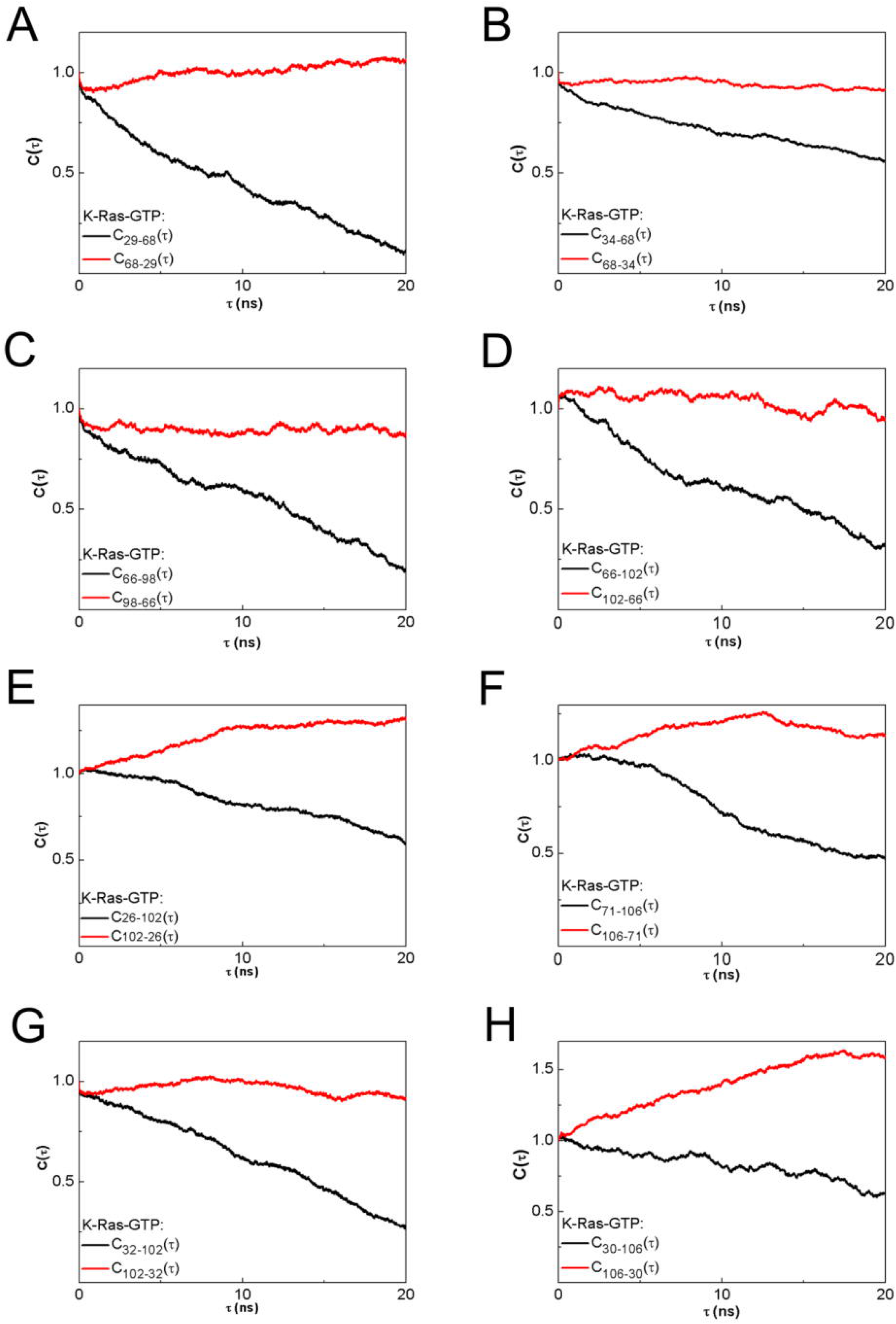
SII fluctuations drive SI fluctuations; α3-L7 motions drive switch region (SI & SII) motions in K-Ras-GTP. Red curves for 〈Δ*R_i_*(*t*)Δ*R_j_*(*t + τ*) show that the fluctuations of residue *i* at time *t* affect the fluctuations of residue *j* at a later time *t*+*τ*. (A) R68 (SII) drives V29(SI). (B) R68 drives P34(SI). (C) E98(α3) drives A66 (α2; SII). (D) R102 (α3) drives A66. (E) S106 (L7) drives Y71 (α2; SII). (F) R102 (α3) drives N26(SI). (G) R102 drives Y32(SI). (H) S106 (L7) drives D30(SI).

Fluctuation decay of K-Ras residues is in the order of 1ns. The red curve persists for time periods that are an order of magnitude longer. The reverse does not show a significant correlation: V29 does not correlate with later fluctuations of R68. Previously, a study reported that SI loop at residues 29–34 swings into the water using V29 and P34 as hinges during Ras inactivation (Noe et al., 2005). We improved on this information by calculating time-delayed correlations and identified that SII residues - especially R68 and D69-sustain active state conformation of SI by driving the motions of hinge residues V29 and P34. Another study also assessed the conformational transition of Ras from inactive to active state (Gorfe et al., 2008), where displacement of SII triggers the active state transition and SI follows SII after a lag time of multiple nanoseconds. Dominance of SII region motions was also observed in several studies (Clausen et al., 2015, Grant et al., 2009a). The nucleotide-bound form behavior is regulated by the relative arrangement of the two switches, rather than their individual conformations. We quantified this by verifying that SI fluctuations follow SII fluctuations in K-Ras-GTP. Since from an information theoretic point of view correlations are regarded as information sources, we conclude that information flows from SII to SI. The directionality originates from the differences in the characteristic decay times. The problem is therefore a problem of dynamics within few nanosecond time periods. Disruption of this flow is expected to interfere with the switch mechanism function, which is the basis of K-Ras activity.

#### α3 and Loop 7 (L7) motions drive switch region (SI & SII) motions in active K-Ras (Figure 6B)

Fluctuations of the helical dimer interface residues of α3, E98 and R102 (Muratcioglu et al., 2015) drive fluctuations of A66 (α2; SII), as shown in Figures 8C and 8D. Additionally, helical dimer interface residue S106 (L7) drives the motion of Y71 (α2; SII) (Figure 8E). On the other hand, fluctuations of R102 (α3) and S106 (L7) drive SI residues N26, D30, Y32 (Figures 8F to 8H).

Correlated motions of α2 and α3-L7 have been described in other studies, which also emphasized the necessity of understanding their effect on protein function (Grant et al., 2009b, Clausen et al., 2015, Grant et al., 2011b). We contribute to this knowledge by identifying their cause and effect relations. Furthermore, in previous studies, starting from the allosteric interaction between α2 and α3-L7, a novel ligand binding pocket, termed p3, which includes residues of L7 was defined and targeted for lead generation (Grant et al., 2011b, Spoerner et al., 2005, Rosnizeck et al., 2010). It was reported that ligand binding to p3 pocket weakens effector protein binding by allosterically stabilizing Ras effector binding site (SI). Another proposed allosteric mechanism is that ligand binding to p3 pocket changes the switch region conformation. Our results implicate the role of allosteric modulation of ligand binding, which may freeze the fluctuations of L7 and stabilize SI motions. This is based on our finding that motions of effector binding site (D30-Y32) are driven by S106 (L7).

#### β2-β3 are both drivers and followers in active K-Ras

Causality calculations suggest the following information flow in fluctuations: ILE21-GLN22 (α1) drives β2-β3 (Figure 6 and S2); which drives Y157 (α5), Q61 (SII) and T74 (SII) (Figure 6 and S3, with details in Table S4). Specifically, the differences in the characteristic decay times in Figures S3A-B demonstrate that information flows from β2-β3 to Y157 (α5). These findings improve on the previous observations of Abankwa et al. where they defined β2–β3 and α5 as a novel conformational switch (Abankwa et al., 2008). Most importantly, we showed that Q61 (SII) motions follow E49 motions (β2-β3) (Figure S3D). Abankwa et al. also observed that mutations in D47-E49 cause hyperactive Ras. Our findings support this too by showing that fluctuations of E49 of the wild type cause fluctuations of the catalytic residue Q61 within SII, whose proper positioning is essential for effective catalysis (Ito et al., 1997). Based on these results, we suggest that mutations in D47-E49 region may alter E49 fluctuations that cause improper Q61 fluctuations. Therefore, GTP catalysis is disrupted which results in constituently active K-Ras.

## Conclusions

We present a novel approach that combines several distinct analysis methods to quantify in detail dynamics of GTP and GDP bound K-Ras, for which a significant amount of experimental and theoretical data already exists in the literature to test our predictions. Oncogenic K-Ras is an extremely high priority drug target in cancer treatment. In order to develop new direct inhibitors that selectively bind to mutant K-Ras conformations while sparing those of WT K-Ras, it is necessary to first understand the dynamic activity of the WT protein in detail. To evaluate the nucleotide binding dependent changes in K-Ras stability, we used stiffness and RMSF calculations and proved that GTP binding rigidifies and hence stabilizes K-Ras motions. These results are in agreement with previous experimental and computational K-Ras studies (Kapoor and Travesset, 2015, Raimondi et al., 2011, Diaz et al., 1995).

Our calculations that use stiffness, RMSF and correlation graphs (Figure 1-2, S1) confirm that GTP binding increases K-Ras stiffness and thereby decreases fluctuation amplitudes, leading to distinct correlation patterns. These striking changes in GTP bound-K-Ras dynamics enable its GTPase activity. Note that this nucleotide exchange is the first step in active to inactive transition (Prakash et al., 2012, Prakash et al., 2015, Zhang et al., 2012, Buhrman et al., 2011, Prakash and Gorfe, 2013, Grant et al., 2011a, Edreira et al., 2009). Overall, our results support the well-established allosteric nature of K-Ras activation (Prakash and Gorfe, 2013), which has been suggested to play an important role in GTPase activity (Grant et al., 2010). Although correlated fluctuations are necessary for allosteric information flow, their longer correlation decay times are also of crucial importance for complete allosteric transition. We calculated time-dependent autocorrelations of fluctuation vectors between residue pairs and discovered that correlations of K-Ras-GTP are stronger and persist for longer correlation times during simulations. Their persistency may allow complete allosteric information flow in K-Ras-GTP.

We broadened our analysis to quantify causality in allosteric regulation of K-Ras function. The most important results from our study are on causality. We applied a simple but powerful method that we defined as *time-delayed correlation* into protein dynamics. To understand K-Ras dynamics, we investigated whether fluctuations of any residue caused fluctuations of another. Our results revealed the information flow in K-Ras switch mechanism and that SII fluctuations drive SI fluctuations. This prediction is an essential validation of our approach, since the dominance of SII motions over SI motions was observed in previous experimental and computational studies (Clausen et al., 2015, Grant et al., 2009a). Surprisingly, in addition to the canonical switch mechanism, our algorithm also revealed causality relations in the novel switch mechanism that includes β2–β3 and α5, where β2–β3 motions drive α5. Moreover, fluctuations of α3-L7 drive fluctuations of SI and SII. Interestingly, previous studies reported that Ras effector binding site (SI) is allosterically stabilized by ligand binding into a novel pocket that includes L7 (Grant et al., 2011b, Spoerner et al., 2005, Rosnizeck et al., 2010). Our results explain the allosteric effect of ligand binding on SI motions by showing the information flow from L7 to SI.

Note that functionally, the identified driver and follower sites do not show an enrichment trend in oncogenic mutations observed in human cancers (from 2266 missense K-Ras mutations observed in all cancers within cBioPortal www.cbioportal.org on July 28, 2016). However, the motions of residue Q61 (SII region), which is the second most frequently mutated residue in cancer patients (113/2266 missense mutations) are driven by those of E49 (β3). While there are no known oncogenic mutations of residue E49, experiments have shown that mutating E49 leads to hyperactive Ras (Abankwa et al., 2008), consistent with what we would expect from a driver of Q61 motions, which also causes the same effect.

Our ongoing research on mutated K-Ras suggests deviations in dynamics from that of WT K-Ras. Any such investigation would first necessitate a deep understanding of the causal relationships in intrinsic K-Ras motions. Understanding intrinsic WT dynamics is imperative and serves as a necessary reference. Our objective behind this detailed analysis is to provide such a reference for any future mutant K-Ras studies. It would be of interest to identify how these findings change when there is a mutation on the protein. Only after identifying these differences can we discover molecules that can eliminate the unfavorable changes caused by the mutations.

The computational tools we introduce in the present work are easily applicable to the analysis of simulation data from different proteins to understand causality in their allosteric regulations which can then be utilized in drug discovery. From this perspective, our approach sets a novel paradigm for drug design that directs attention to changes in protein dynamics. The latter is in close relation to changes in protein function whose restoration to normal is the target of all drug design activities.

## Experimental Procedures

### MD Simulations

We performed all-atom MD simulations for both Mg^+2^GDP- and Mg^+2^GTP-bound K-Ras. As the initial point of all simulations, we used the crystal structure corresponding to PDB ID 4OBE (K-Ras-GDP). For K-Ras-GTP structure, we simply substituted the GDP of K-Ras-GDP structure with GTP using Discovery Studio 4.5 software (BIOVIA, 2015). We solvated each protein in a water box (TIP3 water) where buffering distance between box edges and protein was 12 Å. We applied periodic boundary conditions and added counter-ions to neutralize the system. We used a 2 fs time-step with a 12Å cutoff for Van der Waals interactions and full particle-mesh Ewald electrostatics. We carried out all computations in dynamics procedure for an N, P (1 atm), T (310K) ensemble. We used NAMD 2.10 with AMBER ff99SB and general amber force fields (GAFF). We obtained parameters of GTP and GDP (see Supplemental Experimental Procedures). The initial system energy was first minimized for 10,000 steps, followed by 10,000 steps for equilibration. After equilibration, we performed 300 ns simulations. Atomic coordinates 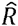 of all atoms were saved every 1 ps. To eliminate all rotational and translational motions, we aligned the trajectories to the initial structure by using VMD software 1.9.2 (Humphrey et al., 1996). We visualized trajectories using VMD.

### Stiffness

We quantified nucleotide-bound K-Ras stiffness using a statistical thermodynamics interpretation of fluctuation correlations (Erman, 2015). We assumed that the interaction between two fluctuating residues *i* and *j* can be represented by a spring, where the spring constant follows from the Gaussian Network Model (GNM) (Haliloglu et al., 1997): 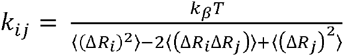 where Δ*R_i_*, is the instantaneous fluctuation of one end of the rod, Δ*R_i_*, is the fluctuation of the other end (Details in Supplementary), *k_B_* is the Boltzmann constant and *T* is the absolute temperature. The spring constant has dimensions of force/length. In GNM spring definition, each residue *i* is attached to *N-1* other residues via *N-1* springs (Haliloglu et al., 1997). Thus, how stiffly a residue *i* is attached to a protein can be quantified by 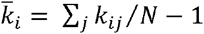 where 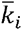 is the mean spring constant for a residue *i*. For stiffness estimates of the complete complexes, we define an overall stiffness parameter *k_overall_* by the expression 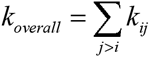. To estimate the stiffness differences in active versus inactive K-Ras, we calculated 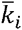 for each residue and *k_overall_* for the protein for both states.

### Stability

We defined the stability of an interacting system of residues as the joint state of reduced RMSF and increased interaction stiffness. RMSF relates to the magnitude of fluctuations of individual residues and stiffness relates to the distance between two residues and therefore they are two independent quantities. (For details see Supplemental Experimental Procedures). A small RMSF and a high stiffness denote increased stability.

### Distance distributions between residue pairs

We calculated the distance between two residues (*i,j*) as 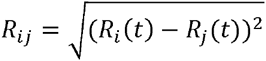. Residue pair distance distributions *W*(*R_ij_*) were calculated by dividing the maximum distance between the pair into small bins and counting the number of observed distances in each bin. All distributions were normalized.

#### Time independent correlations (cross-correlation coefficient map)

Correlations intrinsic to K-Ras structure are defined by the cross-correlation coefficient map, *C*(*ΔR_i_, ΔR_j_*):

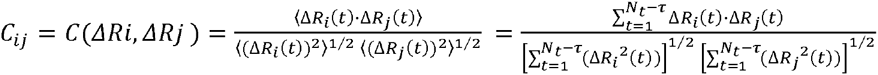

where · denotes the dot product. Correlation varies between -1 and 1. If motions of two atoms are independent, then 〈Δ*R_i_*(*t*) · Δ*R_j_*(*t*)〉 = 0 and *C_ij_* = 0. If the atoms always move in parallel in the same direction, then they are perfectly positively correlated, and *C_ij_* = 1. If they always move in parallel in opposite directions, they are perfectly negatively correlated, and *C_ij_* = -1. Cross-correlation coefficients lie in the range of -1 ≤ *C_ij_* ≤ 1.

### Time delayed correlations, mobility and causality

Time-delayed correlation of two fluctuations is defined by:

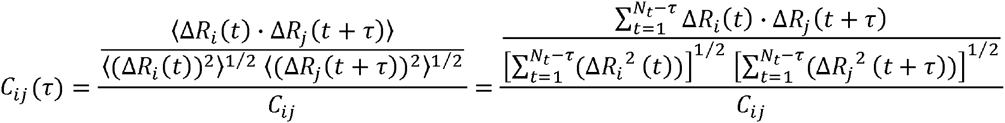

where *C_ij_*(*τ*) represents the correlation of *ΔR_j_* at time *t* + *τ* and earlier values of *ΔR_i_* at time *t*. Similarly, if indices are exchanged, then *C_ij_*(*τ*) represents the correlations of *ΔR_i_* at time *t* + *τ* with earlier values of *ΔR_j_* at time *t*. If the fluctuations of residue *i* drive the fluctuations of residue *j*, then *C_ij_*(*τ*) > *C_ij_*(*τ*). If *C_ij_*(*τ*) > *C_ij_* (*τ*), residue *j* drives residue *i* because the fluctuation *ΔR_j_* at time *t* is correlated with future fluctuations of *ΔR_i_*. However, at *τ*=0, the equality *C_ij_*(0)=*C_ji_*(0) holds.

Note that time-delayed autocorrelation *C_ii_*(*τ*) is the correlation of the trajectory with its own past and future coordinates. If autocorrelation is large, this could correspond to a specific form of “persistence”, a tendency for a system to remain in the same state from one observation to the next.

## Acknowledgements

SV acknowledges 2214-International Doctoral Research Fellowship funding from The Scientific and Technological Research Council of Turkey (TUBITAK). ZHG and SV were supported by the LUNGevity Foundation and the start-up funds to ZHG from Icahn School of Medicine at Mount Sinai.

